# Steady state visual evoked potentials reveal a signature of the pitch-size crossmodal association in visual cortex

**DOI:** 10.1101/2022.11.07.515442

**Authors:** Placido Sciortino, Christoph Kayser

## Abstract

Crossmodal correspondences describe our tendency to associate sensory features from different modalities with each other, such as the pitch of a sound with the size of a visual object. While such crossmodal correspondences (or associations) are described in many behavioural studies their neurophysiological correlates remain unclear. Under the current working model of multisensory perception both a low- and a high-level account seem plausible. That is, the neurophysiological processes shaping these associations could commence in low-level sensory regions, or may predominantly emerge in high-level association regions of semantic and object identification networks. We exploited steady-state visual evoked potentials (SSVEP) to directly probe this question, focusing on the associations between pitch and the visual features of size, hue or chromatic saturation. We found that SSVEPs over occipital regions are sensitive to the congruency between pitch and size, and a source analysis pointed to an origin around primary visual cortices. We speculate that this signature of the pitch-size association in low-level visual cortices reflects the successful pairing of congruent visual and acoustic object properties and may contribute to establishing causal relations between multisensory objects.

## Introduction

Humans often associate sensory features from different modalities with each other (Kohler, 1929, 1947; Spence, 2011; Spence and Deroy, 2013; Evans and Treisman, 2010; Fort and Schwarz, 2022). For example, a low-pitched sound is typically associated with the larger of two visual objects. These crossmodal associations have been defined as the intuitive pairing between perceptual attributes of stimuli in different senses and are known also as ‘‘crossmodal similarities”, “crossmodal correspondences'’, ‘‘synesthetic associations’’ or ‘‘sound symbolic associations’’. Crossmodal associations have been reported for various combinations of sensory modalities and features, and have been studied in a variety of experimental paradigms that range from verbal reports to speeded classification tasks (Spence, 2011). Still, these behavioural results are only partly paralleled by neuroimaging data and the cerebral origins of crossmodal associations remain unclear.

In particular, it remains debated whether crossmodal associations are mediated by low-level sensory cortices or whether they are the result of high-level integration processes in semantic or object identification networks in associative cortices (McCormick et al., 2021, 2018). Under the current working model of multisensory perception, both a low- and a high-level account seem plausible. In this model, multisensory representations emerge along multiple stages of the sensory pathways, commencing in low-level sensory regions with automatic interactions constrained by stimulus location or timing and continuing with higher specificity in association regions that establish the weighted combination of sensory modalities depending on their congruency and task relevance (Bizley et al., 2016; Cao et al., 2019; Noppeney, 2021; Rohe and Noppeney, 2015; Shams and Beierholm, 2022).

In one previous study, participants were trained on specific sound-symbolic associations and EEG recordings revealed association-specific activations around 140 ms post-stimulus onset, letting the authors speculate about an involvement of low-level sensory regions (Kovic et al., 2010). Such an account would be in line with object-related multisensory congruency effects at even earlier latencies (Molholm et al., 2004) and the activation of primary auditory and visual cortices in the Bouba-Kiki effect (Peiffer-Smadja and Cohen, 2019) or other multisensory paradigms (Brang et al., 2022; Kayser et al., 2010; Lakatos et al., 2009; Schroeder and Foxe, 2005). However, EEG studies on the pitch-size association pointed to effects at longer latencies, possibly arising from parietal and frontal regions (Bien et al., 2012; Stekelenburg and Keetels, 2016).

A high-level account is also supported by fMRI studies that reported neural correlates of crossmodal associations in higher associative areas such as parietal, temporal and frontal areas or pointed to a more distributed neural network involving in multisensory attention, memory- or imagination-related processes (McCormick et al., 2018, 2021). However, multisensory attention and imagination have neurophysiological correlates that can extend or propagate back into low-level sensory regions (Ferrari and Noppeney, 2021; Talsma et al., 2010; Vetter et al., 2014; Zuanazzi and Noppeney, 2020), which again fuels speculations that crossmodal associations may commence or at least shape neurophysiological processes at early states of cortical processing (Peiffer-Smadja and Cohen, 2019).

We here directly tested the hypothesis that associations between auditory and visual features are reflected in activity from low-level visual cortices - hence that their signatures are visible in functionally defined low-level sensory regions. To this end, we exploited visual steady-state evoked potentials (SSVEP) as markers of neural activity arising from low-level visual cortices (Norcia et al., 2015; Regan, 1966; Vialatte et al., 2010). The SSVEP is elicited by the rapid presentation of visual stimuli at a fixed rate and has been used to study the involvement of visual cortices in processes such as figure ground segregation, perceptual dominance, binocular rivalry, attention, object recognition and memory (Alpers et al., 2005; Appelbaum et al., 2008, 2006; Brown and Norcia, 1997; Kaspar et al., 2010; Müller and Hübner, 2002; Silberstein et al., 2001; Sutoyo and Srinivasan, 2009; Walter et al., 2012). We chose this paradigm as it comes with two specific advantages: first, it relies on a functionally-induced and well-studied neurophysiological marker of sensory-specific activations (Di Russo et al., 2007; Lauritzen et al., 2010; Norcia et al., 2015; Tsoneva et al., 2021; Vialatte et al., 2010). Second, it allows probing the stimulus-selective response of visual cortex to one of two simultaneously presented stimuli, hence emphasising the ‘binding’ nature of multisensory associations. This binding effect may be characteristic of crossmodal correspondences but is more difficult to assess in paradigms involving only one stimulus per modality (Evans and Treisman, 2010; Parise and Spence, 2009).

In our experiment, we focused on three putative crossmodal associations of pitch with the visual features size, hue and chromatic saturation. All three features reflect basic visual attributes that are encoded in visual cortices and hence their association with pitch may have a correlate in low-level sensory regions. An association between pitch and size has been reported in multiple behavioural studies, whereas the associations of pitch with hue and pitch with saturation have been found less consistently (Anikin and Johansson, 2019; Bernstein et al., 1971; Caivano, 1994; Giannakis, 2001; Hamilton-Fletcher et al., 2017; Melara, 1989; Moos et al., 2014; Orlandatou, 2012; Ward et al., 2006). The comparison of these three putative associations allowed us to probe whether they potentially all share a similar neurophysiological correlate. We tested these associations by presenting two concurrent flickering visual stimuli that differed along one specific feature dimension (size, hue or saturation) together with one sound (low or high in pitch) or in the absence of a sound. Thereby we probed both the influence of the presence of any sound on the visual cortical response and the selective influence of low or high-pitched sounds on the response to specific sizes, hues or saturation levels. Our results reveal a congruency-related effect for the pitch-size association in both the SSVEP and behavioural data, but not for the other two associations. Given that the SSVEP mainly originates from low-level visual cortices (Di Russo et al., 2007; Lauritzen et al., 2010) and that the congruency effect in the present data was source localized to visual cortex, this speaks for a correlate of the pitch-size association in low-level sensory regions and shows that SSVEP can be exploited to study crossmodal processes.

## Methods

### Participants

A total of 26 participants (16 females, mean age = 27.5) participated in the experiment. The sample size is comparable to previous studies using the SSVEP procedure and testing crossmodal associations in behaviour or brain activity (Appelbaum et al., 2006; Kaspar et al., 2010; Lauritzen et al., 2010; Parise and Spence, 2012, 2009). All participants gave written informed consent prior to participation and all procedures were approved by the Ethics Committee of Bielefeld University. Participants were compensated for their time (10 Euro/hour). The experiments were performed in a darkened and electrically shielded room (E:box, Desone). Participants sat on a comfortable chair in front of a computer screen with a grey background. The data from one participant had to be excluded (see below), hence all data are for a sample of 25 participants.

### General paradigm

The paradigm was designed to probe three audio-visual associations: between pitch and the size of a visual object, between pitch and hue, and between pitch and chromatic saturation. We chose the association between size and pitch as this has been reported robustly across many studies (Evans and Treisman, 2010; Gallace and Spence, 2006; Parise and Spence, 2008, 2009). The associations between pitch and the hue or saturation of visual stimuli have been reported before, but the robustness of these remains debated (Anikin and Johansson, 2019; Bernstein et al., 1971; Caivano, 1994; Hamilton-Fletcher et al., 2017; Melara, 1989; Moos et al., 2014; Simpson et al.,1956; Ward et al., 2006). Concerning hue, we focused on blue and yellow, as these are considered safer than for example red and green when presented as flickering stimuli (Drew et al., 2001; Tello et al., 2015).

Visual stimulation was based on an SSVEP procedure (Burkitt et al., 2000; Di Russo et al., 2007; Lauritzen et al., 2010; Norcia et al., 2015; Vanegas et al., 2013; Vialatte et al., 2010). During each trial two visual stimuli were presented for 6.2 seconds, one flickering at 7.5 Hz and one at 12 Hz, presented on a uniform grey background (15 cd/m^2^). These specific frequencies were chosen as they are incommensurate and in the range of those used in most SSVEP studies probing the encoding of visual features, image content or attention (Appelbaum et al., 2008, 2006; Brown and Norcia, 1997; Kaspar et al., 2010; Norcia et al., 2015; Silberstein et al., 2001; Vialatte et al., 2010; Walter et al., 2012). The two stimuli were placed 11 degrees of the left and right of a central fixation point. The individual stimuli were defined by their shape (irrelevant for the specific hypotheses tested here), their size, hue and saturation (Figure 1). During each trial the two visual stimuli differed by only one feature, which defined the visual feature whose association with the sound was probed in the respective trial. The specific shapes were a triangle, a circle and a square. The shape was varied in order to minimise intramodal associations between the manipulated features and other properties such as angularity, spikiness, or the number of edges (Dreksler and Spence, 2019). The small and large visual stimuli subtended about 3.5 and 6.5 degrees of visual angle in diameter. For the specific values of hue and saturation see below. During each trial, either one of two sounds was presented (high pitch: 1000 Hz tone, low pitch: 500 Hz tone) or no sound was present. Across trials, the different combinations of visual features, flicker frequency, side of presentation and auditory conditions were counterbalanced.

**Figure 1.**
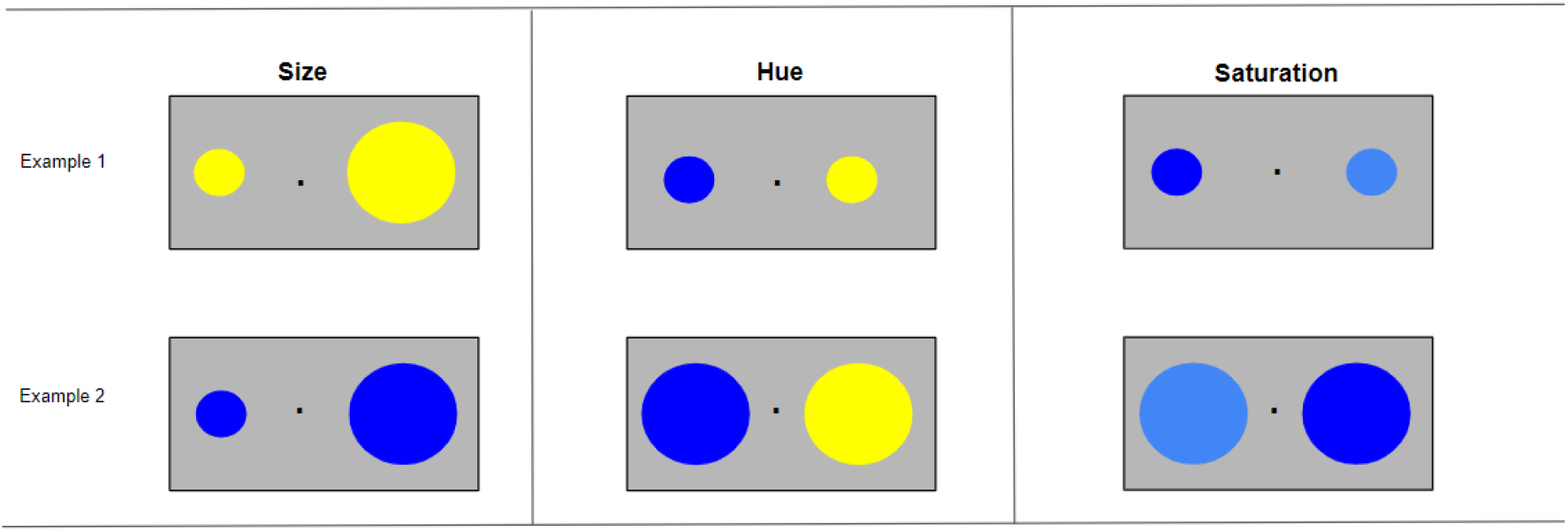
Schematic of the stimuli used to test the associations of pitch with size, hue and saturation. The two visual stimuli presented on each trial differed by one dimension (either hue, size, saturation) and could take one of three shapes (circle, square, triangle) to avoid biasing the results by using a single shape. The two visual stimuli flickered independently at 7.26 and 11.6 Hz for a period of 6.2s and were accompanied by either a high pitched tone (1000 Hz), a low pitched tone (500 Hz) or no sound The individual features, flicker frequencies, sound conditions and side of presentation were counterbalanced across trials.

As a result, each trial either probed one specific crossmodal association (pitch-size, pitch-hue or pitch-saturation) or presented the visual stimuli in the absence of an auditory stimulus. When a tone was presented, one of the two visual stimuli was congruent with the sound according to the associations tested, the other was incongruent. Congruencies were operationally defined as follows and similar to the previous literature. Congruent conditions were high pitch & small size and low pitch & large size; high pitch & yellow hue and low pitch & blue hue; high pitch & high saturation and low pitch & low saturation. A total of 288 trials were presented, divided into 6 blocks. Each association was tested in 96 trials each that were divided into two blocks. Each trial started with a fixation period (600 to 900 ms, uniform), followed by the 6.2 s stimulation period. Inter-trial intervals lasted 1100 to 1400ms (uniform).

At the end of the stimulation period one of the two shapes was presented for an additional 4 frames (33.3ms) and hence was visible longer than the other. Participants were asked to indicate which of the two shapes was presented longer by pressing the matching right or left arrow key on a keyboard. Participants were instructed to be as fast and as accurate as possible. Importantly, on trials with a sound present the visual stimulus remaining longer could be either the congruent or incongruent one with the concurrent pitch. We employed this task to ensure that participants maintained attention to both visual stimuli and to probe for a behavioural correlate of the crossmodal association.

Stimuli were presented using the Psychtoolbox (3.0.15) from Matlab (R2017a). Visual stimuli were presented on an Asus #PG279Q monitor running at 120Hz refresh rate, with stimuli at 7.5 Hz being presented each 16th frame and those at 12Hz each 10th frame. Each stimulus was drawn for two consecutive frames on the screen. The timing of the visual stimuli was verified for each trial and participant by inspecting the time stamps provided by Psychtoolbox and was generally verified using an oscilloscope. This revealed that the actual screen refresh rate was 116.9 ± 0.12 Hz (mean ±s.e.m. across experimental days) rather than the nominal 120Hz. Hence, the two actual flicker frequencies were effectively 7.26 and 11.6 Hz. These values were very consistent across participants and were used for subsequent data analysis. The auditory stimuli were presented at 62 dB SPL from two speakers positioned to the left and right of the monitor. Sound intensity was calibrated using a sound level-meter (Bruel & Kjaer 2250).

### Selection of hue and saturation levels

Because the perceived hue and saturation of chromatic stimuli is subjective we used participant-wise calibrated stimuli that were subjectively perceived as ‘yellow’ and ‘blue’ and had similar perceived lightness and saturation against a uniform grey background. We considered this important as otherwise two visual stimuli may also differ in their subjective saliency. For this purpose we conducted a pre-test. In a first step participants were presented a HSV colour wheel (saturation and lightness values of 0.7) on a grey background and were instructed to select the “most pure blue” or “most pure yellow” direction respectively using a mouse. Subsequently, participants were presented with one of the selected hues (at fixed saturation) along with a coloured bar presenting the other hue in varying degrees of saturation. They were then asked to select the level of saturation of the second hue matching that of the first. In a third step they were asked to equalise the level of lightness in a similar manner. This iterative process was randomized and repeated 4 times (twice matching yellow against blue and twice blue against yellow). We then calculated the average of each HSV attribute and used these for each participant as hue, saturation and lightness. While lightness was fixed through the experiment for each hue, saturation varied between high- and low-saturation trials. High saturation was defined as the average of the chosen saturation values, while low saturation was defined by dividing the chosen high value by two.

### EEG recording and pre-processing

EEG signals were acquired using a 128 channel BioSemi (BioSemi, B.V.) system with Ag-AgCl electrodes mounted on an elastic cap (BioSemi). Four additional electrodes were placed at the outer canthi and below the eyes to obtain the electro-oculogram (EOG). Data were acquired at 1028 Hz sampling rate and electrodes offset were kept below 25 mV. Data analysis was performed using MATLAB (The MathWorks Inc.; R 2017A) and the FieldTrip toolbox (ver. 22/09/2019)(Oostenveld et al., 2011).

Data pre-processing was similar to our previous work (Kayser and Kayser, 2020; Park and Kayser, 2021; Sciortino and Kayser, 2022a, 2022b). The data were band-pass filtered between 0.6 and 90 Hz and resampled to 200 Hz. Electrodes with faulty contacts were interpolated (on average 0.65 ± 0.15 per participant, mean ± s.e.m.). Subsequently the data were denoised using Independent component analysis (ICA). We identified and removed components reflecting oculomotor or muscle activity following previous literature and using a semi-automatic procedure (Campos Viola et al., 2009; Hipp and Siegel, 2013; O’Beirne and Patuzzi, 1999; Sciortino and Kayser, 2022a, 2022b). In total we removed 11.5 ± 0.10 mean ± s.e.m. components per participant. To extract the SSVEP the pre-processed data were epoched around the stimulation interval, removing the first 500 ms following stimulus onset and omitting the last frames of the stimulus that introduced the behavioural task.

### Quantification of the SSVEP response

To visualise and quantify the SSVEP we computed the power spectra of the data in Fieldtrip. For this we resampled the data to 40 Hz and computed the time-averaged power for individual trials using Hanning windows. These power estimates were obtained for each electrode separately and were then averaged across either all trials and conditions, or only for specific conditions of interest (see below). We visually inspected these spectra and confirmed that they reflect the expected peaks at the two stimulation frequencies (Figure 2A). The data from one participant clearly deviated from this expected pattern and the data from this participant were excluded.

**Figure 2.**
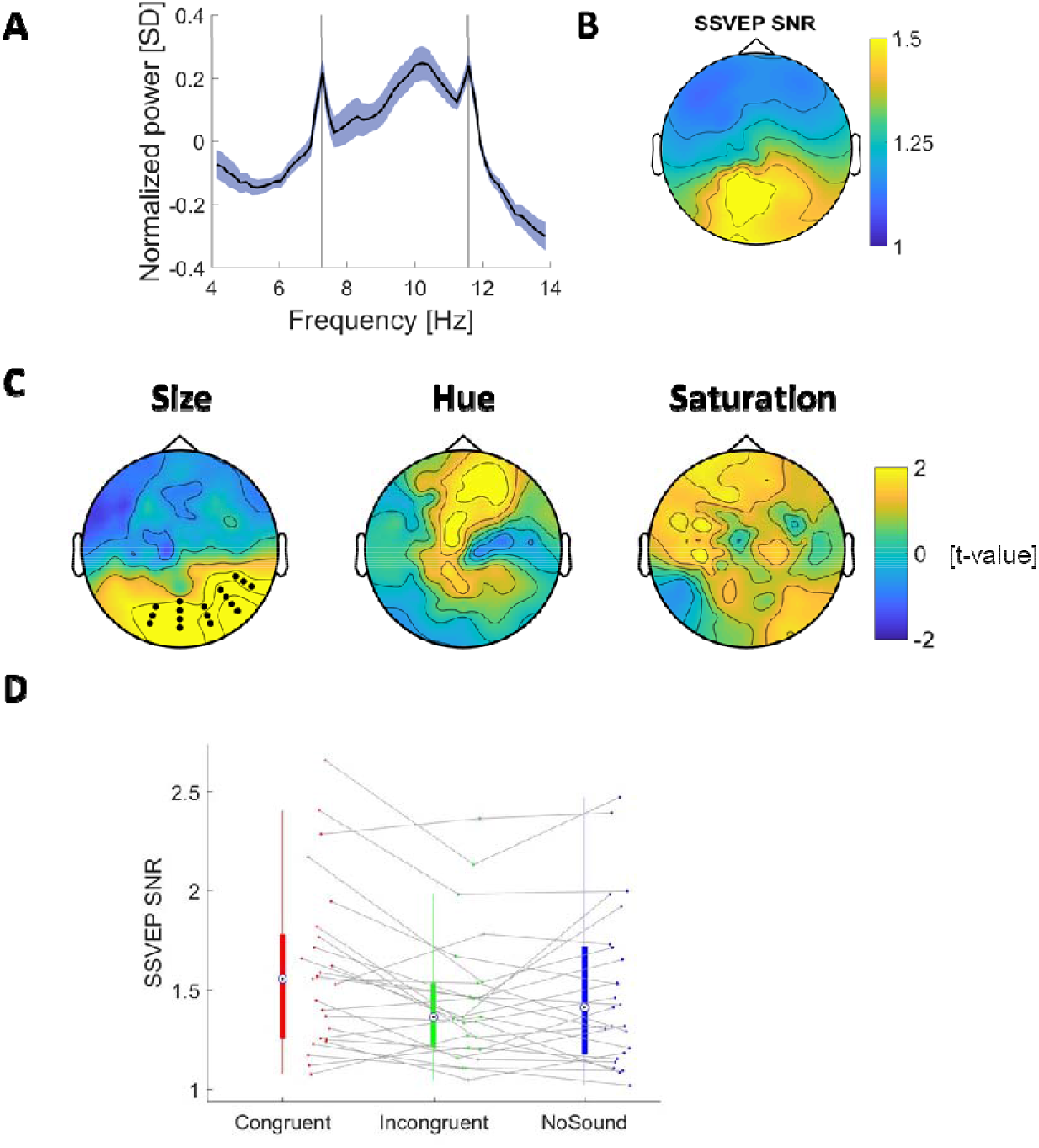
SSVEP response and crossmodal associations. **A)** Normalized EEG spectra during SSVEP stimulation for occipital electrodes. For visualization the log power spectra were normalized within each participant (z-scored and linearly detrended to remove the 1/f dependency). The graph shows the group mean (n=25) and standard error. The two lines indicate the SSVEP stimulation frequencies. **B)** Topographies of the group-average SSVEP SNR, obtained across both stimulation frequencies and all conditions. The SNR quantifies the power at each stimulation frequency against the average of the power in immediately neighbouring frequency bins. **C)** Topographies of the electrode-wise t-maps for a difference between congruent minus incongruent conditions for each association. For pitch-size we found a significantly higher SSVEP response for congruent conditions over occipital electrodes (p=0.02, cluster-based permutation test). **D)** SSVEP responses for individual conditions and participants in the pitch-size association obtained in the significant cluster.

Based on the stimulation paradigm we expected two local peaks in these spectra around 7.26 and 11.6Hz. Because the shape of the EEG spectra and their overall power differs between participants, we computed the signal to noise ratio at these two stimulation frequencies (SNR) as a participant-wise normalised measure of the SSVEP response (Lopez-Gordo et al., 2011; Vialatte et al., 2010). For this we divided the power near each stimulation frequency (7.26 ± 0.1 Hz and 11.65 ± 0.1 Hz respectively) by the average power in neighbouring bands. These neighbouring bands were defined as −0.8Hz to −0.3Hz below and +0.3Hz to +0.8Hz around each stimulation band. This SNR based approach has the advantage that the power of interest and the noise estimates are obtained from the very same data epochs, and hence does not require the additional assumption that a putative baseline period and the stimulus period are comparable in terms of sensory or cognitive factors. A group-level topographical visualisation of the SSVEP SNR is shown in Figure 2B. This reveals the expected strong SSVEP-SNR over occipital electrodes, indicating that the stimulation frequencies adopted were successful in generating a strong signal in visual cortices (Di Russo et al., 2007; Lauritzen et al., 2010; Norcia et al., 2015; Vanegas et al., 2013; Vialatte et al., 2010).

To probe our main question, we computed this SNR for specific constellations of trials. First, we split trials according to the sound conditions (no sound, high or low pitch). In addition, by design of our paradigm, we split trials according to whether the congruent (incongruent) visual stimulus on each trial was presented at the lower or higher flicker frequency. This allowed us to selectively compute the SSVEP response for trials in which the sound was congruent (incongruent) with the stimulus at each stimulation frequency. Note that by design each trial enters this analysis twice, once via the congruent and one via the incongruent visual stimulus. We then computed the SSVEP SNR across all congruent (incongruent) trials at each frequency separately and averaged the results across stimulation frequencies. This was done as our main question pertains to the effect of sound (presence or congruency) and we did not expect cross-modal associations to induce frequency specific effects in the SSVEP. Rather, any robust crossmodal association effect should emerge independently of stimulation frequency. This analysis resulted for each participant in one SNR for each crossmodal association, congruency condition (or no sound) and electrode.

### Source analysis

We implemented a confirmatory source analysis using the DICS beamformer in FieldTrip (Gross et al., 2001; Oostenveld et al., 2011). For this we relied on a standardized head model based on the average template brain of the Montreal Neurological Institute as single participant MRI data were not available. Lead-fields were computed using a 3D grid with 6mm spacing. We localized the trial- and participant-averaged condition wise power at frequencies of interest. Specifically, to derive the SSVEP SNR we localized the power at the two stimulation frequencies of interest and the power in the side-bands used to compute the SNR separately. We then computed the SNR in source space for each stimulation frequency and averaged these. Figure 3 shows the overall SNR and the congruency effects for pitch and size.

**Figure 3.**
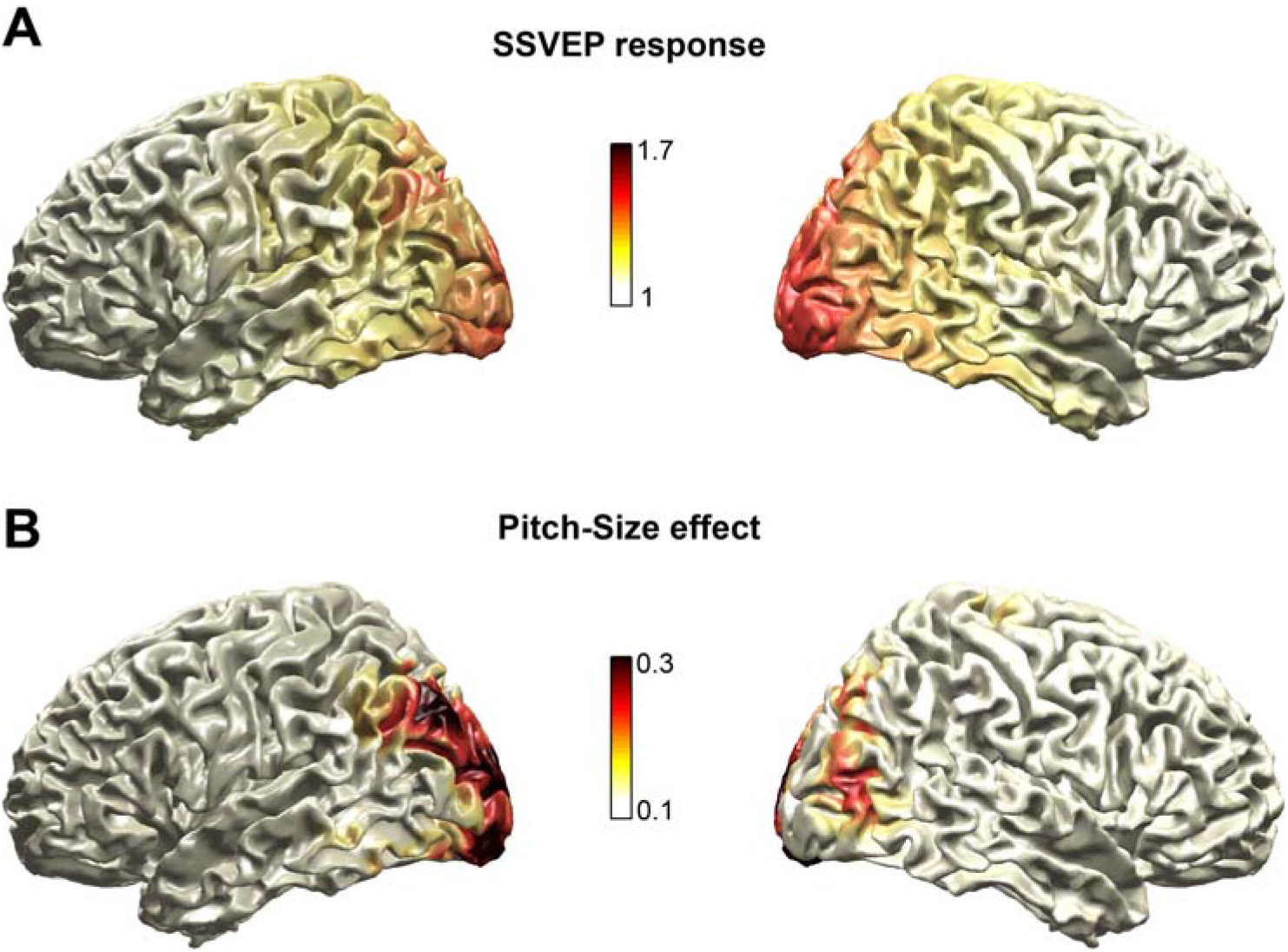
Source results for the pitch-size association. **A)** Source projection of the group-averaged SSVEP response (SNR), across both stimulation frequencies and both congruent and incongruent epochs. **B)** Pitch-size congruency difference in source space, with positive numbers reflecting a stronger source response for congruent stimuli.

### Statistical analysis

For the EEG data the comparisons of interest concerned the group-level SSVEP responses between congruent and incongruent pairs. This contrast was derived for each of the three associations. To test for a difference while correcting for multiple tests across electrodes we relied on a cluster-based permutation procedure (Maris and Oostenveld, 2007). We computed electrode-wise paired t-tests between congruent and incongruent epochs and thresholded these at a first-level threshold of p<0.01 (two-sided). Significant electrode-clusters were aggregated using the cluster-mass (using a minimal cluster-size of 3 electrodes) and the cluster-wise statistics in the actual data were compared to a surrogate distribution obtained from 5000 randomizations. For each significant cluster we report the p-value and the cluster-mass as test-statistics. As effect size we report Cohen’s D at the electrode with maximal effect sizes. We also implemented a post-hoc analysis within the significant cluster obtained for the pitch-size association (c.f. Fig. 2D). Here we for example probed whether the SSVEP response was affected by the general presence of a sound (vs. the no-sound condition). For this we first averaged the SNR across electrodes and then contrasted this between conditions using paired t-tests.

To compare the mean response times between conditions (congruent vs. incongruent, or against no-sound), we first averaged the log-transformed response times across trials, removing RTs faster than 150 and slower than 1500 ms as outliers. We then contrasted conditions using a Wilcoxon sign rank test. Effect sizes were obtained using Point biserial correlation (denoted rrb).

## Results

### SSVEP reveals an effect of association-congruency

The SSVEP stimulation induced robust responses over occipital electrodes (Figure 2A,B). These were quantified as the SNR of the induced power at the two stimulation frequencies relative to the power in immediately neighbouring frequency bands. Figure 2B shows the group-level SNR topography, which reveals the expected peak over occipital electrodes.

We asked whether this SSVEP response differed between congruent and incongruent conditions, hence whether the visual cortical response to a given stimulus is affected by the pitch of the tone presented at the same time. This statistical comparison was implemented for each association, correcting for multiple comparisons using a cluster-based permutation procedure. For the association between pitch and size we found a significantly stronger SSVEP response for the congruent condition (n = 25 participants; p=0.02, tsum = 49.1, Cohen’s D = 0.93, 17 electrodes; Fig. 2C). This pitch-size congruency difference was strongest over occipital electrodes, in line with a potential origin in visual cortex. We verified that this congruency effect held also for the two individual stimulation frequencies, and for both was the SNR stronger for congruency trials, although the effect size was weaker as for the analysis combining both stimulation frequencies (Cohen’s D of 2.62 and 3.2 for 7.2 and 11.6 Hz respectively). For the other two associations, pitch-hue and pitch-saturation, we found no significant differences in SSVEP responses between conditions (the maximal effect sizes across electrodes were as follows: pitch-hue D_max_ =0.46, D_min_ = −0.20; pitch-saturation D_max_ =0.42, D_min_ = −0.11).

### Source results

We implemented a source analysis to visualize the sources underlying the pitch-size congruency effect. Figure 3A shows the SSVEP response, which is strongest over bilateral occipital regions and confirms the expected origin of the SSVEP response in visual cortex. Figure 3B shows the pitch-size congruency effect, in analogy to Figure 2C. This pitch-size congruency difference was strongest over occipital regions, with a broad peak centered on MNI coordinates [−6, −100, 10] and covering the AAL atlas regions ‘Calcarine’ and ‘Occ Sup’ on both hemispheres (Tzourio-Mazoyer et al., 2002).

### Sounds do not facilitate visual responses in general

Given that many studies have shown generic multisensory benefits for neurophysiological responses in low-level regions we also compared the SSVEP response between trials in which a sound was present compared to trials without sound. For this we focused on the cluster of electrodes revealing a significant congruency-effect in the pitch-size association (Fig. 2D). We contrasted the SSVEP response between the no-sound trials and each individual condition and with the sound conditions combined (average of congruent and incongruent) (Fig. 2D). While this analysis confirmed the known congruency effect (t=3.3, p=0.0025, Cohen’s D=0.67) it revealed no significant differences between no-sound trials and congruent (t=1.97, p=0.06, D=0.39) or incongruent trials (t=-1.4, p=0.17, D=0.28) or between sound and no-sound trials (t=0.15, p=0.87, D=0.03). This suggests that the SSVEP response was not generally affected by the presence of a sound but rather points to a specific effect of the pitch-size congruency.

### Response times

As a behavioural measure of the crossmodal correspondences we included a speeded detection of a deviant stimulus at the end of the stimulation period. Specifically, one of the two stimuli stayed on longer for a brief moment (33.3 ms) and participants had to indicate which one (left or right). We reasoned that the congruency between the sound, which was already present for the 6 seconds prior to this task, may interact with the detection of this event for the congruent visual stimulus. We hence compared congruent and incongruent trials for each association (Figure 4).

**Figure 4.**
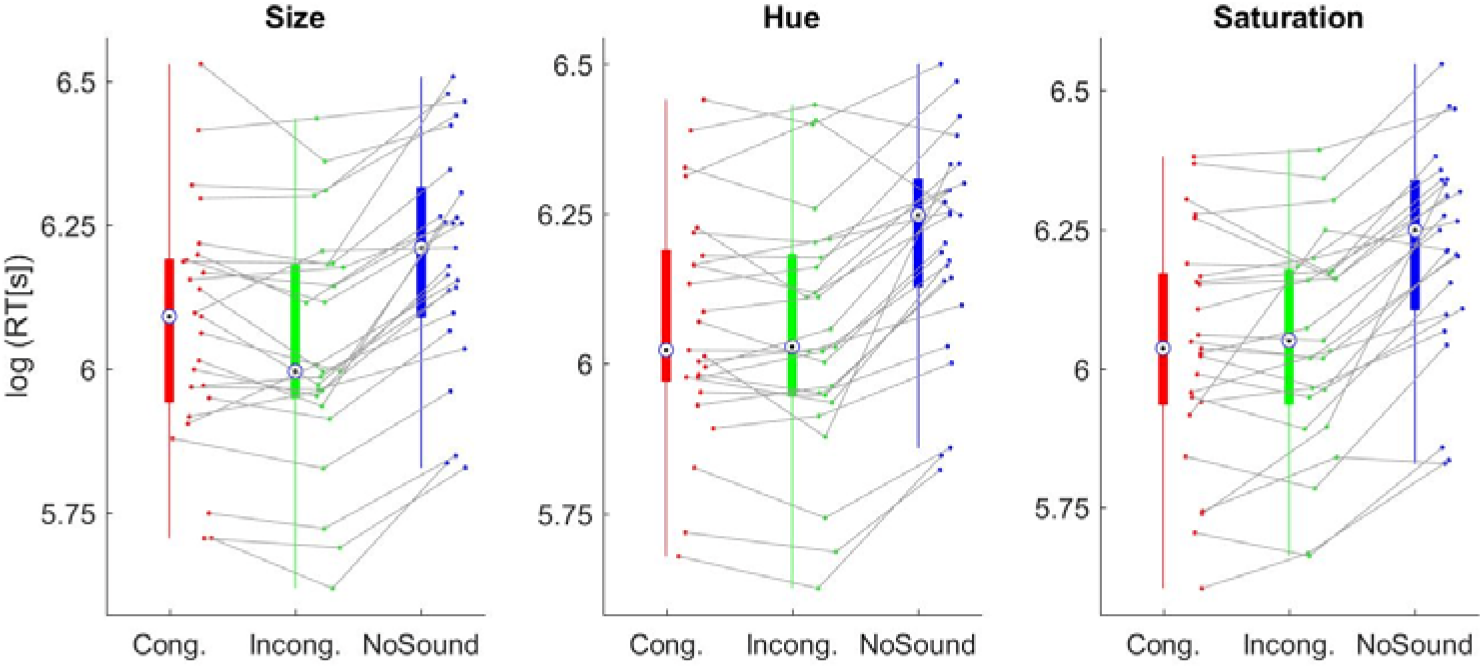
Behavioural response times. The graphs show the log-transformed response times for the three conditions and each of the association blocks. Dots indicate individual participants (n = 25).

For the pitch-size association responses in the incongruent trials were significantly faster compared to congruent trials (n=25, Wilcoxon sign-rank test, Z=2.25, p = 0.02, rrb =0.51). No significant results were found for the other associations (Hue: Z=1.06, p = 0.29, rrb =0.24; Saturation: Z=-0.39, p = 0.69, rrb=-0.08). We also probed for a general facilitation of reaction times by the presence of any sound, contrasting response times on trials with a sound and those without sound. This revealed that participants were significantly faster when a sound was present (Size= Z=4.34, p < 0.01, rrb =0.99; Hue: Z=4.05, p < 0.01, rrb =0.93; Saturation: Z=4.37, p < 0.01, rrb =1). Hence in the behavioural data we observed a general facilitation of a sound on visual task performance as well as a congruency effect for the pitch-size association. However, this behavioural effect was not significantly correlated with the congruency effect in the SSVEP across participants (p=0.18, r=-0.27).

## Discussion

Humans consistently associate properties of stimuli in different sensory modalities with each other (Spence, 2011; Spence and Deroy, 2013; Spence and Sathian, 2020). Well-known examples are the association between pitch and size or between phonological features and visual shapes such as in the Bouba-Kiki effect (Kohler 1929, 1947; Fort and Schwartz, 2022). In the latter the word Bouba is associated with the round and Kiki with the more spiky among two visual shapes. Despite considerable work on multisensory perception in general, the neurophysiological correlates of such associations remain debated (Spence and Sathian, 2020). While some studies have argued for an involvement of low-level sensory regions others have implied high-level processes in association cortices (Bien et al., 2012; Kovic et al., 2010; McCormick et al., 2021, 2018; Peiffer-Smadja and Cohen, 2019; Stekelenburg and Keetels, 2016) and recent principled attempts to localize the underlying correlates using fMRI have not provided clear results (McCormick et al., 2021, 2018). To address this question, we hence opted for an approach that has received little attention in multisensory studies, but which comes with specific benefits: measuring SSVEP’s we set out to test the hypothesis that neurophysiological correlates of audio-visual associations are visible in activity arising from functionally defined low-level visual cortices. In support of this, we found a pitch-size congruency effect that was strongest over occipital electrodes and in primary visual cortical source regions. In contrast, the associations between pitch and hue or saturation did not reveal significant neurophysiological or behavioural effects, suggesting that these are less robust than the pitch-size association, as we discuss below.

### Robust associations of pitch with size but not hue or saturation

The association between the pitch of a sound and the size of a visual object possibly arises from the physical properties of the objects surrounding us, where larger animals or objects produce lower frequency sounds compared to smaller ones (Bee et al., 2000; Carello et al., 1998; Tecumseh Fitch and Reby, 2001). This association has been reported in numerous studies using implicit behavioural measures such as reaction times or explicit matching responses (Bien et al., 2012; Evans and Treisman, 2010; Mondloch and Maurer, 2004; Parise and Spence, 2012).Our data support the robustness of this association both at the behavioural and neurophysiological level.

In contrast, the associations between pitch and chromatic properties such as a specific hue or saturation level have been reported in some but not in other studies, suggesting they are less prevalent in the population or weak in terms of their behavioural effect sizes (Anikin and Johansson, 2019; Bernstein et al., 1971; Caivano, 1994; Hamilton-Fletcher et al., 2017; Melara, 1989; Moos et al., 2014; Simpson et. al, 1956; Ward et al., 2006). Furthermore, these studies disagree about which specific chromatic features were associated with a sound. For example, some reported the association of high pitch with yellow or green relative to blue or purple (Simpson et al., 1956; Moos et al., 2014; Giannakis 2001, Orlandatou 2012), while others suggested that the specific pitch-hue or pitch-saturation associations may depend on the precise values of acoustic pitch and visual features used (Anikin and Johansson, 2019; Hamilton-Fletcher et al., 2017; Wrembel, 2009) or the context of other sound frequencies presented in the same experiment (Ward et al., 2006). Importantly, some studies did not control for the subjectively perceived lightness when manipulating hue or saturation (de Thornley Head, 2006; Simpson et a., 1956; Ward et al., 2006; but see Anikin and Johansson, 2019), which may be crucial given that an association between pitch and lightness has been reported as well (Brunel et al., 2015; Hubbard, 1996; Marks, 1987, 1975, 1974). Hence, some of the discrepancies in the previous literature may be a result of subjective differences of the respective stimuli along multiple chromatic dimensions. To minimise such confounding factors, we relied on stimuli that were calibrated on a participant-level for their subjective hue, saturation and lightness. Based on these stimuli, we did not find significant congruency effects for pitch-hue or pitch-saturation, neither in the behavioural nor in the neurophysiological data. Hence, our results suggest that these associations may be not as robust as the association between pitch and size, or may indeed depend on the precise nature of the specific stimuli or of the behavioural paradigm employed, as some have speculated (Hamilton-Fletcher et al., 2017). For sure, more work is required to better understand the nature of sound-chromatic associations and their prevalence.

### Correlates of cross-modal associations in low-level sensory cortices

We introduced an SSVEP based paradigm to probe crossmodal associations for two reasons. First, given the known origin of the SSVEP response in visual cortex it allows establishing whether or not a neurophysiological correlate from these sensory regions relates to crossmodal associations (Burkitt et al., 2000; Di Russo et al., 2007; Lauritzen et al., 2010; Norcia et al., 2015; Vanegas et al., 2013; Vialatte et al., 2010). Second, it allowed us to probe the response to simultaneously presented congruent and incongruent visual stimuli with the same sound, and therefore directly taps into the characteristic binding property of crossmodal associations (Parise and Spence, 2009).

Our results reveal a stronger SSVEP response occipital regions for the visual stimulus whose size was congruent with the pitch. This provides direct evidence for a low-level neurophysiological correlate of the pitch-size association, in which the visual response to a visual stimulus matching a concurrent sound in some feature-specific manner is enhanced (Kovic et al., 2010; Molholm et al., 2004; Peiffer-Smadja and Cohen, 2019). Such an interpretation is in line with previous speculations suggesting a low-level or short-latency correlate of object-related multisensory congruency effects in general (Brang et al., 2022; Kayser et al., 2017; Kovic et al., 2010; Molholm et al., 2004; Peiffer-Smadja and Cohen, 2019; Sadaghiani et al., 2009) and in line with the working model of multisensory processing. This model proposes that interactions in low-level sensory regions are governed by low-level congruencies of the multiple inputs, such as their temporal synchronization or spatial alignment, while multisensory integration in association cortices is more functionally and task-specific (Bizley et al., 2016; Cao et al., 2019; Noppeney, 2021; Rohe and Noppeney, 2015). Our results show that these multisensory processes in low-level visual cortices are also sensitive to congruencies pertaining to basic features such as size and pitch. Given that these features are correlated in everyday life based on the physical attributes of the objects surrounding us (Spence, 2011; Parise et al., 2014), this suggests that early crossmodal interactions may be adapted to our natural environment. In fact, crossmodal associations may reflect the binding of multisensory features often occurring together, possibly to aid the process of multisensory causal inference. This process reflects the brain’s decision as to whether or which sensory features should be combined during perception and which not to (Körding et al., 2007; Noppeney, 2021). While studies have attributed a central role of frontal regions in this process (Cao et al., 2019; Ferrari and Noppeney, 2021; Rohe and Noppeney, 2015), the consequences of causal inference may be fed back to low-level sensory regions to support the grouping of matching sensory features there. This speculation is also supported by our finding that the congruency effect in the SSVEP response was not accompanied by a general enhancement from the presence of an acoustic stimulus per se. This suggests that the association-related auditory influence on visual cortex is distinct and more specific than the previously reported general modulation of visual responses by acoustic inputs (Brang et al., 2022; Lakatos et al., 2009; Murray et al., 2016; Petro et al., 2017; Wang et al., 2008).

Our data cannot disambiguate the contribution of feed-forward and feed-back processes given that the SSVEP response was elicited and analyzed over a longer stimulation interval and given that the SSVEP originates from a number of low-level visual areas and spreads to other visual areas as well (Norcia et al., 2015; Tsoneva et al., 2021; Vanegas et al., 2013; Vialatte et al., 2010). Hence, it is possible that the neurophysiological signature visible in occipital regions was generated in response to multisensory feedback, for example as a result of causal inference, but possibly also including processes such as attention or memory. In fact, in our paradigm the sound may have facilitated the perceptual binding of the auditory input with the congruent of the two visual shapes, for example via the spread of attention across modalities (Talsma et al., 2010). This attention-spread may be one factor contributing to the enhanced SSVEP response, which is known to be sensitive to attention effects in general (Appelbaum and Norcia, 2009; Lauritzen et al., 2010; Norcia et al., 2015). An alternative account could be predictive coding, whereby auditory regions relay information that is predictive of concurrent or expected features to visual cortices, possibly via direct cortico-cortical connections (Falchier et al., 2002; Garner and Keller, 2022; Morrell, 1972; Petro et al., 2017). Hence, while our results demonstrate the existence of a neurophysiological signature of cross-modal correspondences in sensory cortices, future work still needs to uncover the specific functional and computational benefits.

For the SSVEP paradigm, we opted for frequencies in the alpha range, which are known to emphasize the involvement of low-level visual cortices, including primary areas (Di Russo et al., 2007; Lauritzen et al., 2010; Tsoneva et al., 2021; Vanegas et al., 2013). Still, the specific generators of the SSVEP response may depend also on the specific stimulation frequency and the activations evoked by the SSVEP may travel along the visual pathway (Alonso-Prieto et al., 2013; Appelbaum and Norcia, 2009; Cottereau et al., 2011; Liu-Shuang et al., 2014). However, we found that both the SSVEP responses itself and the pitch-size congruency effect are strongest over occipital electrodes and were source localized with primary visual cortices, all in direct support of an origin in low-level visual cortical regions.

We found signatures of the pitch-size association in both the SSVEP and the behavioural data. Yet, the SSVEP response to congruent visual stimuli was enhanced, while the response times were longer when the deviant stimulus was congruent with the sound, and the two effects were not significantly correlated across participants. While these two signatures seemingly have opposing directions we see reasons for why this may be plausible. First, the SSVEP response was obtained over a 6 second stimulation period. In contrast, the behavioural response was obtained only at the very end of the stimulation period, hence after prolonged exposure to the stimuli. Such prolonged stimulation may induce adaptation effect which can dissociate sensory responses and behaviour (Blakemore and Campbell, 1969; Blakemore and Sutton, 1969; Gibson and Radner, 1937; Leopold et al., 2005). Second, the stronger SSVEP responses for congruent visual stimuli may reflect the enhanced binding of these specific visual features with the sound. Binding the congruent visual object with the sound may foster its perceptual constancy or persistence, which may then be detrimental for the detection of a change in the congruent (bound) compared to the incongruent (not bound) object. For example, participants have higher thresholds for discriminating spatial or temporal conflicts in synesthetically matched compared to mismatched audio-visual stimuli (Parise and Spence, 2009). Multisensory binding may impede or slow down the neurophysiological access to the individual audio and visual sensory components, therefore in our case reducing the behavioural detection of changes relative to the congruent pair (Ernst and Banks, 2002; Ernst, 2006). Third, other studies measuring SSVEPs also found opposing effects on SSVEP response strength and behaviour. In general, the SSVEP response provides an index of visual cortical neural responses. However, since the EEG reflects pre- and post-synaptic responses this does not only include the mere processing of sensory input but reflects the aggregate effect of all intracortical processes. For example, opposing effects between SSVEP and behavioural performance have been found for several tasks such as working memory, spatial attention and object recognition (Kaspar et al., 2010; Silberstein et al., 2001; Walter et al., 2012) suggesting that an apparent mismatch between SSVEP signatures and behavioural data may be a more general phenomenon.

### Unspecific multisensory benefits for behaviour

Our behavioural data also speak in favour of a general facilitation of visual task performance by the presence of a sound compared to the no-sound condition. Similar sound-induced benefits of visual perception have been reported for task-irrelevant or uninformative sounds in a range of paradigms (Fiebelkorn et al., 2011; Frassinetti et al., 2002; Huang et al., 2011; Jaekl and Soto-Faraco, 2010; Lippert et al., 2007). The present data corroborate the existence of such effects. We did not see a general sound-modulation of the SSVEP responses, which speaks for a different neurophysiological origin of these unspecific effects compared to the cross-modal association. However, unspecific multisensory effects were also attributed to early sensory cortices (Brang et al., 2022; Kayser et al., 2009; Mercier and Cappe, 2020; Pérez-Bellido et al., 2022). Hence, it is possible that both association-related and unspecific multisensory processes influence low-level sensory responses but do so by different pathways and neural mechanisms and give rise to distinct neurophysiological signatures in the EEG.

### General conclusion

Despite their prominence in behavioural and psychophysical studies, the neurophysiological mechanisms underlying cross-modal associations remain unclear. Using an SSVEP stimulation protocol known to engage low-level visual cortices we show that the association between pitch and the size of a visual object influences activity in these visual regions. Hence our data support a low-level account of crossmodal associations.

